# Amyloid plaque deposition accelerates tau propagation via activation of microglia in a humanized APP mouse model

**DOI:** 10.1101/2020.09.22.308015

**Authors:** Kevin A. Clayton, Jean Christophe Delpech, Shawn Herron, Naotoshi Iwahara, Takashi Saito, Takaomi C. Saido, Seiko Ikezu, Tsuneya Ikezu

## Abstract

Microglia have an emerging role in development of tau pathology after amyloid plaque deposition in Alzheimer’s disease, although it has not been definitively shown. We hypothesize that plaque-associated activated microglia accelerate tau propagation via enhanced phagocytosis and secretion of tau. Here we show that the injection of adeno-associated virus expressing P301L tau mutant into the medial entorhinal cortex (MEC) in humanized APP^NL-G-F^ knock-in mice induces exacerbated tau propagation in the dentate gyrus compared to wild type mice. Depletion of microglia dramatically reduces accumulation of phosphorylated tau (pTau) in the dentate gyrus as well as an extracellular vesicle (EV) marker, Tumor susceptibility gene 101, co-localized in microglia. Mac2^+^ activated microglia secrete significantly more EVs compared to Mac2^−^ microglia in APP^NL-G-F^ mice *in vivo* when injected with lentivirus expressing EV reporter gene mEmerald-CD9, suggesting enhanced EV secretion by microglial activation. Our findings indicate that amyloid plaque-mediated acceleration of tau propagation is dependent on activated microglia, which show enhanced EV secretion *in vivo*.

## Introduction

Microglia are the immune cells of the central nervous system and possess a number of specific roles including synaptic pruning [1, 2], release of pro-inflammatory and anti-inflammatory cytokines [3, 4], as well as surveying for and phagocytosing pathologic insults [5–7]. Recently, a class of disease-activated microglia unique to neurodegeneration (referred to as “MGnD” or “DAM) has been characterized along with their role in the alleviation or exacerbation of neurodegenerative disorders [8, 9]. MGnD exhibit characteristics similar to activated microglia and possess a unique molecular signature that is regulated by Triggering Receptor Expressed on Myeloid cells 2 (TREM2) and ApoE, which disrupts maintenance of CNS homeostasis [8, 9]. MGnD microglia, which are identified via immunofluorescence against markers such as C-type lectin domain family 7 member A (Clec7A) and Galectin-3 (Mac2), reside around amyloid plaques and phagocytose not only aggregated proteins, but also apoptotic neurons and synapses. There is still ongoing discussion as to whether MGnD are ultimately beneficial or harmful in neurodegenerative disease. MGnD may play a key role bridging the amyloid plaque toxicity and the tau pathology development in Alzheimer’s disease (AD). Amyloid plaques precede tau pathology in AD and are believed to initiate or build upon mechanisms responsible for tau pathology. Indeed, previous reports showed that amyloid-beta pathology accelerates tau pathology development in different mouse models [10–12]. Lately, the interplay between the spread of pathological tau and neuritic plaques as determined by hyperphosphorylated tau (p-tau) has been a subject of investigation as pTau^+^ neuritic plaques, referred to as “NP Tau”, appears to affect sequestration and spread of pathologic tau seeds in a manner that may be dependent on microglia [10, 13]. It is still undermined whether microglia play a critical role for the accelerated tau propagation in the presence of Aβ plaque deposition.

One of the methods in which we can assess the effect of microglia on AD pathology is through their selective depletion. This is accomplished in models of conditional knockdown of microglia, targeted depletion, or via administration of Colony stimulating factor 1 receptor (CSF1R) inhibitors [14–18]. Using CSF1R inhibitors along with other means of selective microglia depletion, researchers examined the effect of depletion on amyloid plaque deposition, tau pathology and spread, synaptic integrity, as well as cognition in a variety of mouse models of neurodegeneration [19–23]. We previously established a rapid tau propagation model in which AAV-P301L Tau is injected into the medial entorhinal cortex (MEC) where tau is expressed and eventually propagated to the granular cell layer (GCL) of the dentate gyrus (DG) in a manner facilitated by microglia and exosomes, small extracellular vesicles (EVs) synthesized in multivesicular bodies [22]. In the present study, we investigated how microglia may facilitate tau propagation following activation by amyloid plaques by injecting AAV-P301L Tau into the MEC of APP^NL-G-F^ knock in mice, which develop robust amyloid plaque formation by endogenously expressing 3 APP mutations [24].

We revealed that amyloid burden accelerated tau propagation in APP^NL-G-F^ mice compared to wild type (WT) mice and depleting microglia dramatically reduced tau propagation to the GCL in both cases. Additionally, we observed loss of Clec7A^+^ microglia as well as an exosomal marker, Tumor susceptibility gene 101, around amyloid plaques. Increased deposits of NP Tau and amyloid plaque following microglia depletion in APP^NL-G-F^ mice further suggest MGnD actively phagocytose protein aggregates and propagate them through subsequent exosome secretion. This posits a possible mechanism to explain enhancement of tau propagation with amyloid toxicity and amyloid associated microglia. Lastly, we constructed a novel lentivirus to express CD9 conjugated to mEmerald green fluorescent protein (mE-CD9) in a microglia-specific manner, and quantified the release of mEmerald-CD9^+^ EV particles at single cell resolution *in vivo*. We found that the degree of EV release was over 3 times higher from Mac2^+^ MGnD compared to Mac2^−^ microglia. These findings provide additional evidence that microglia are important for transferring misfolded and aggregated tau between cells via EVs and suggest that amyloid deposition may enhance tau propagation through MGnD induction of microglia.

## Results

### Microglia Depletion increases plaque deposition

In order to examine the effect of microglia depletion on plaque pathology and tau propagation, APP^NL-G-F^ (homozygote for the *App* loci with *APP*^NL-L-G-F^) and WT mice aged 4 months were fed chow containing the CSF1R inhibitor (PLX5622) or placebo for 4 weeks prior to AAV-Tau P301L injection and for an additional month following AAV injection until euthanasia (Fig 1A). We didn’t observe any adverse effect of PLX5622 administration during the treatment period. Immunofluorescence against P2RY12, a chemoreceptor highly and uniquely expressed in microglia [25], showed that P2RY12 was almost completely absent (>92%) after over 8 weeks of drug treatment in both groups (Fig 1B-C). In the AD brain, microglia become activated in response to amyloid pathology and migrate to the region to compact and phagocytose Aβ plaques [26–29]. We hypothesize that microglial depletion will alter the Aβ plaque compaction in APP^NL-G-F^ mice. Circularity of thioflavin-S^+^ plaques was dramatically reduced by microglia depletion, suggesting that Aβ plaques were less compacted in the absence of microglia (Fig 1D-E). Consistent with this data, overall Aβ plaque positive area, number and size were significantly increased by microglial depletion (Fig 1E). 3D surface renderings of thioflavin-S plaques using IMARIS revealed a dramatic decrease in sphericity and a dramatic increase of plaque volume and area (Fig S1A-B).

**Figure 1.**
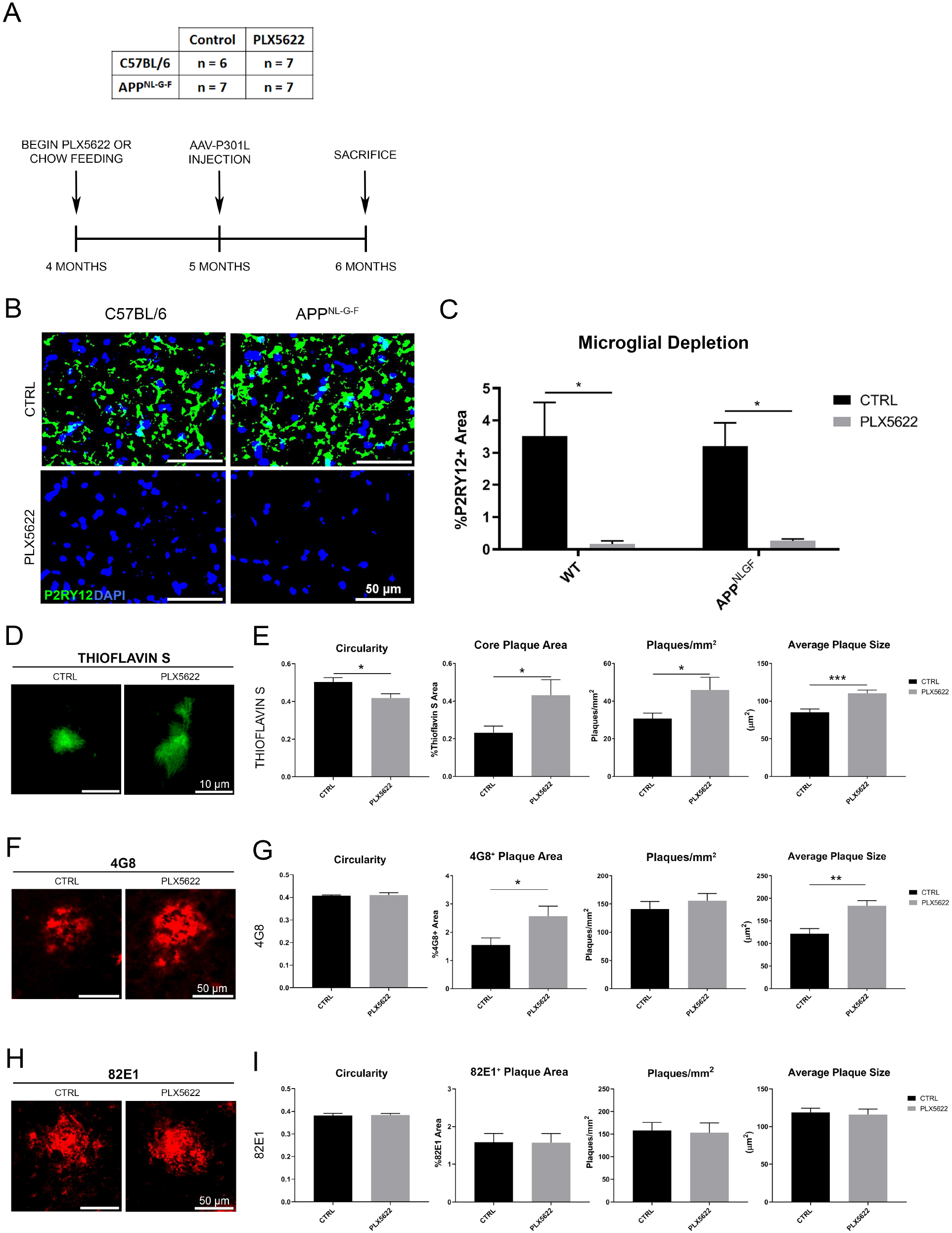
PLX5622 treatment ablates microglia and increases amyloid burden. **A.** Schematic of study design with PLX5622 administration. The table displays the number of mice used per experimental group. **B.** Representative images of P2RY12 staining in the cortical region from WT and APP^NL-G-F^ having received control (CTRL) or PLX5622 chow. **C.** Unbiased quantification of percentage P2RY12^+^ area in the cortex. **D.** Representative images of Thioflavin-S staining in the cortex. **E.** Unbiased quantification of plaque characteristics in Thioflavin-S stained slices in the cortex. **F.** Representative images of 4G8 staining in the cortex. **G.** Unbiased quantification of plaque characteristics in 4G8 stained slices in the cortex. **H.** Representative images of 82E1 staining in the cortex. **I.** Unbiased quantification of the plaque characteristics in 82E1 stained slices in the cortex. All values displayed in **A-I** represent the the mean ± standard error (SEM) for a minimum of 6 animals per group. Graphs comparing values across all 4 groups were analyzed via 2-way analysis of variance (ANOVA) with Tukey post-hoc analysis for individual comparisons. Graphs comparing two groups were analyzed via Unpaired t-test. \**p* < 0.05, \*\**p* < 0.01, \*\*\**p* < 0.001, between indicated groups.

These findings were mostly reproduced in diffuse amyloid staining with 4G8 (detecting Aβ17-24, Fig 1F-G), specifically in the size and number of plaques. Interestingly, immunofluorescence for diffuse amyloid with an alternate antibody, 82E1, revealed no differences in plaque pathology after microglia depletion in the same brains (Fig 1H-I). Moreover, 82E1 possessed a distinct staining pattern from 4G8 (Fig S1C). The 82E1 antibody detects Aβ1-16, which is analogous to the 6E10 antibody, is most commonly used in studies assessing the effect of microglia depletion on amyloid burden (Supplementary Table S1). The discrepancy in the Aβ plaque detection between 82E1 and 4G8 antibodies may be an important consideration for future studies. In addition, immunofluorescence against Glial fibrillary acidic protein (GFAP) was also used to assess the impact of microglia depletion on astrogliosis (Fig S1D). In addition, immunofluorescence against glial fibrillary acidic protein (GFAP), a marker for activated astrocytes, was also used to assess the impact of microglia depletion on astrogliosis (Fig S1D). There was a robust increase in astrogliosis around Aβ plaques in APP^NL-G-F^ mice compared to WT mice that was unaffected by PLX5622 treatment (Fig S1D-E). Together, these results strongly suggest that microglia play a significant role in compaction and number of Aβ plaques in APP^NL-G-F^ mice at 4 to 6 months of age.

### Enhanced tau propagation and its reduction by CSF1R inhibition in APP^NL-G-F^ mice

Abnormally phosphorylated tau (pTau) propagates from the MEC to the hippocampus the anatomically connected regions as seen in the process of tau pathology development in AD. The perforant pathway describes the tri-synaptic circuit of neuronal connections from the MEC to the dentate gyrus (DG), Cornus Ammonis 3 (CA3) and CA1 (Fig 2A) [30]. To determine the effect of microglial depletion on tau propagation via the perforant path in APP^NL-G-F^ mice, animals were treated 4 weeks with PLX5622 or control chow from 4 months of age. AAV2/6 pseudotyped synapsin-1 promoter-driven transgene expression of P301L MAPT mutant (AAV-P301Ltau) was injected into the MEC (AP: 4.75, ML: 2.90, DV: 4.64) in APP^NL-G-F^ and WT mice (Fig. 2A) at 5 months of age as previously described (Fig. 1A) [22], and the animals were fed with either PLX5622 or control chow for 4 weeks until the end point of the study. Tau propagation from the MEC to the DG was assessed by the immunofluorescence against p-tau (AT8, detecting pSer202/pSer205 tau). Following injection of AAV-P301L tau, we observed strong AT8^+^ cell soma staining in the MEC and in the GCL of the DG in APP^NL-G-F^ mice and also in WT mice to a lesser extent (Fig 2B). Furthermore, we also found AT8^+^ cells in the Hilus and CA1 regions in both WT and APP^NL-G-F^ mice (Fig S2A-B), suggesting that pTau was propagated through not only the perforant pathway, but also possibly the temporoammonic pathway, which projects from EC layer II to CA1 (Fig 2A). This result was consistent with previous reports showing an augmented effect of amyloid plaques on tau propagation [10, 12, 31]. Strikingly, microglial depletion effectively suppressed the tau propagation in both groups (Fig 2C), and the inhibitory effect of microglia depletion on tau propagation was more prominent in APP^NL-G-F^ compared to WT mice (Fig 2C).

**Figure 2.**
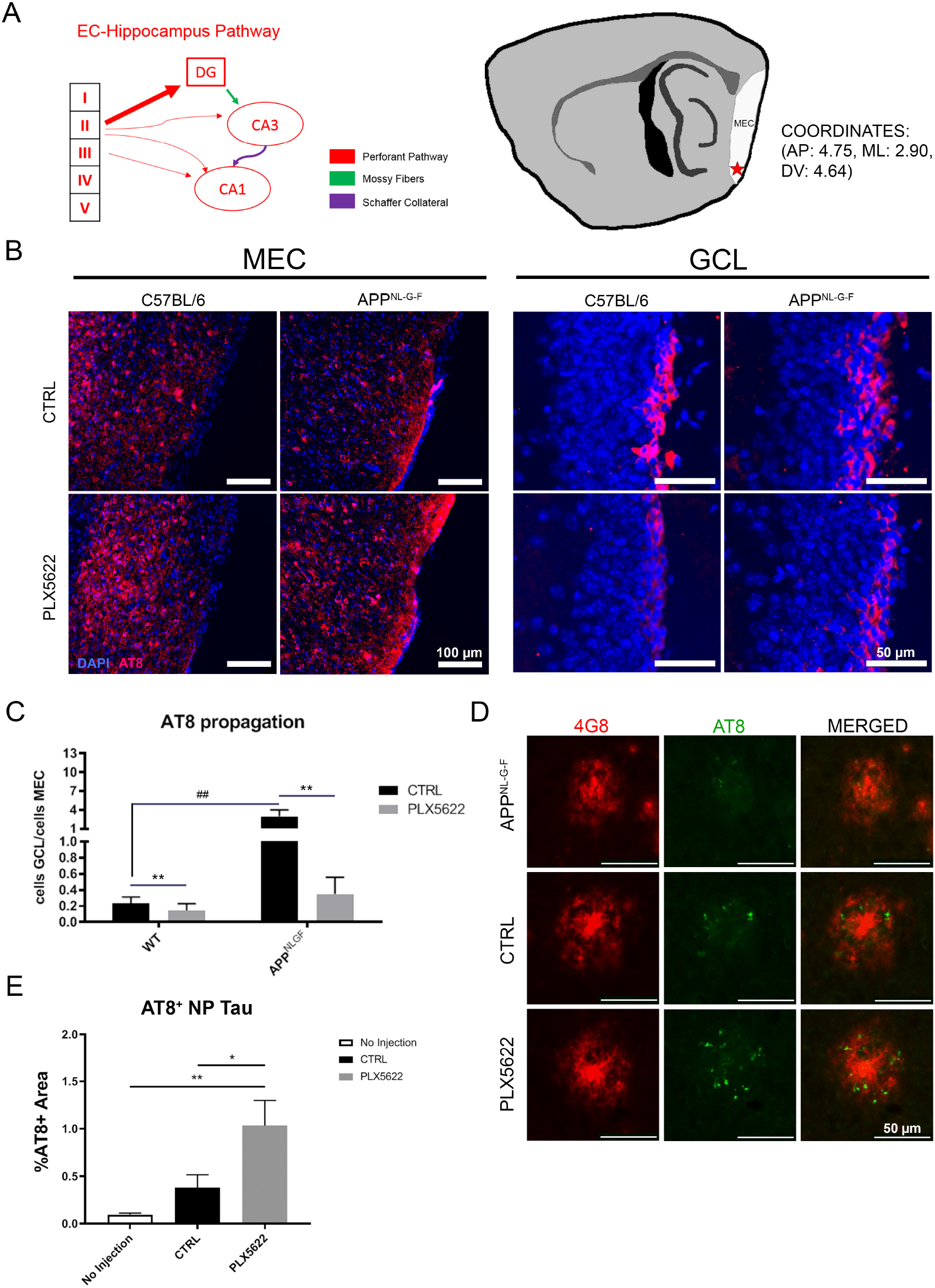
Aβ deposition accelerate tau propagation in microglia-dependent manner, while microglial depletion enhances NP tau formation. **A.** Schematic of injection coordinate and tri-synaptic pathway. **B.** Representative images of AT8 staining in the MEC and the GCL region of the DG. **C.** Unbiased quantification of AT8^+^ or HT7^+^ cell propagation from the MEC to the GCL. Tau propagation values across all 4 treatment groups are displayed. **D.** Representative image of plaques in the MEC of non-injected APP^NL-G-F^ mice, injected mice receiving control chow, and mice receiving PLX5622. **E.** Unbiased quantification of percentage AT8^+^ NP tau within the MEC. All values displayed in **A-E** represent the mean ± SEM. Graphs comparing values across 4 groups were analyzed via 2-way ANOVA with Tukey post-hoc analysis for individual comparisons. Graphs comparing values across 3 groups were analyzed via 1-way ANOVA with Fisher’s LSD post-hoc analysis for individual comparisons. * *p* < 0.05, ** *p* < 0.01, ##*p* < 0.01 between WT-CTRL and APP^NL-G-F^-CTRL.

In addition to AT8^+^ neurons, AT8^+^ plaque tau deposits were investigated. One recent study demonstrated the significant effect of TREM2 mutation on NP tau formation in tau fibril-injected APPPS1-21 mouse brains, suggesting the role of microglia in NP tau development [13]. We thus examined the effect of AAV-P301Atau-mediated tau propagation on NP tau formation in APP^NL-G-F^ mice with or without microglial depletion. Interestingly, AT8^+^ NP tau surrounding plaques were increased following AAV-P301L tau incubation in the MEC, which was further significantly enhanced by microglial depletion (Fig 2D-E). This suggests that tau expressed in MEC can also spread to NP tau, which are actively phagocytosed by plaque-associated microglia. Together, these data provide strong evidence that Aβ deposition exacerbates tau propagation, plaque-associated MGnD microglia play an essential role on transferring tau from the MEC to the DG, and they may also actively phagocytose NP tau in AAV-P301Ltau-injected APP^NL-G-F^ mice.

### Plaque-associated activated microglia secrete C1q and EVs

C1q is a protein complex known to be part of the complement system and is also mostly derived from microglia [32]. It is highly deposited on synapses in the molecular layer of the DG [33, 34]. Expression level of C1q in the outer molecular layer was significantly reduced in APP^NL-G-F^ mice following microglia depletion, revealing the absence of microglia-derived proteins (Fig 3A-B). In the AD brain, microglia become activated in response to amyloid pathology and migrate to the region to compact and phagocytose plaque material [26–29]. We next examined the activated microglia around plaque regions by Clec7A staining. Clec7A^+^ MGnD microglia were detected around the plaques in APP^NL-G-F^ mice, and were ~80% eliminated following PLX5622 treatment (Fig 3C-D). Microglia produce and secrete exosomes more efficiently following activation [35]. We found striking co-expression of Tsg101 in Clec7A+ MGnD microglia, which were also diminished by PLX5622 treatment (Fig. 3C, E), suggesting that plaque-associated MGnD microglia secrete (EVs). This is also supported by the data from Clec7A^+^ microglia isolated from 6 months-old APP^NL-G-F^ mice, which show significantly upregulated gene expression of EV makers, such as *Cd9* and *Cd63*, along with MGnD maker *Apoe* (Fig 3F). Clec7a+/-were isolated via fluorescence-activated cell sorting for markers Ly6C, CD11b, FCLRS, and Clec7a (Fig S3A). These findings suggest MGnD hyper-secrete EVs in APP^NL-G-F^ mice.

**Figure 3.**
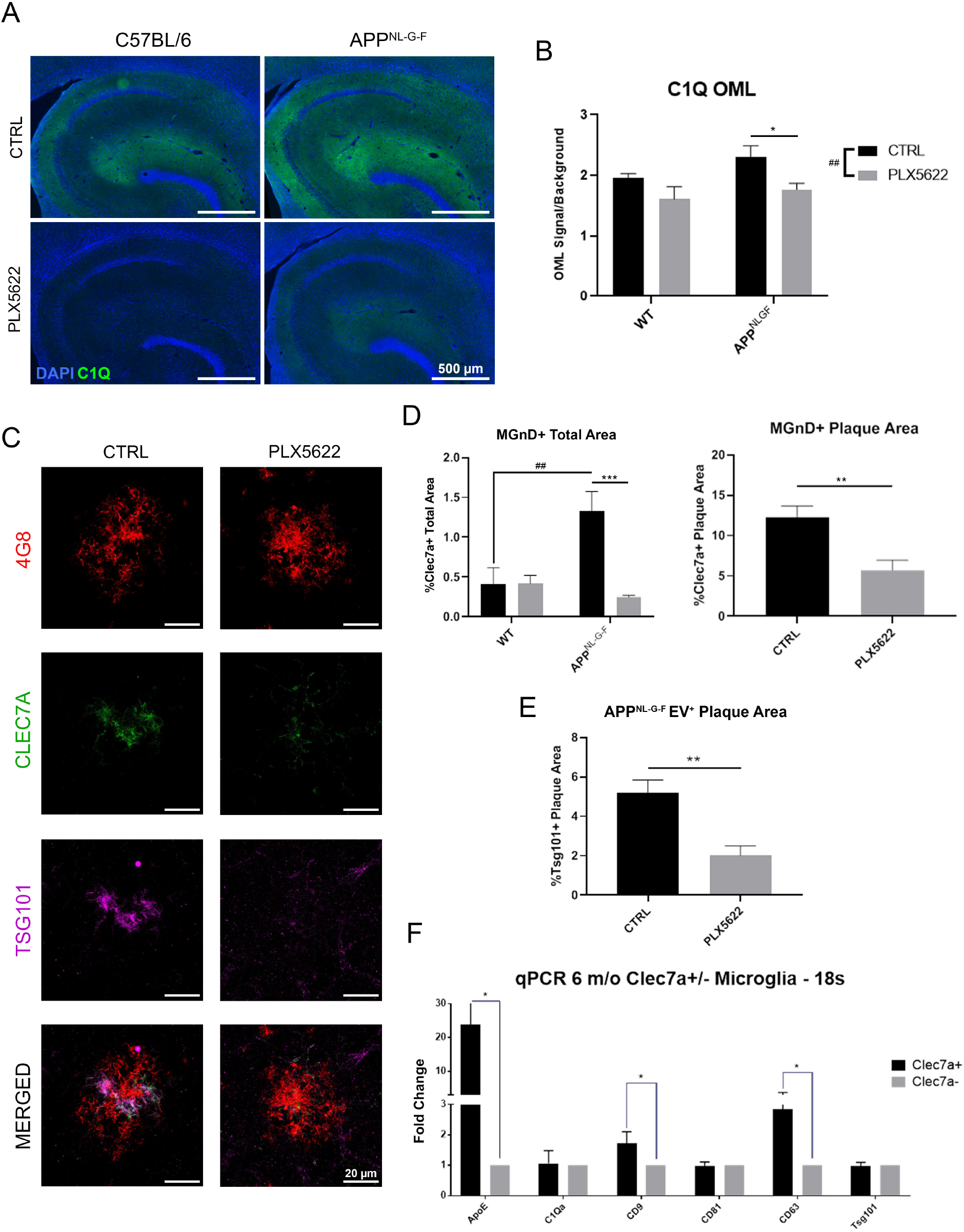
Plaque-associated microglia are responsible for secretion of C1q and EVs. **A.** Representative images of C1Q + DAPI staining in the hippocampal region. **B.** Unbiased quantification of C1Q intensity in the OML of the hippocampus. **C.** Representative stacked confocal images of 4G8, Clec7A, and Tsg101 staining of plaques in the OML. **D.** Unbiased quantification of percentage Clec7A^+^ area across the OML for all groups and of the plaque-specific regions. **E.** Unbiased quantification of Tsg101 intensity of plaque-positive regions in the OML compared to background signal. **F.** qPCR data displaying fold-increases in EV-associated markers Cd81, Cd63, Cd9, Tsg101, and C1qa expressed in Clec7A^+^ over Clec7A^−^ microglia. All values displayed in **A-F** represent the mean ± SEM for a minimum of 3 animals per group. Graphs comparing values across all 4 groups were analyzed via 2-way ANOVA with Tukey post-hoc analysis for individual comparisons. Graphs comparing two groups were analyzed via Unpaired t-test. **\*\*** *p* < 0.01, **\*\*\*** *p* < 0.001, **\*\*\*\*** *p* < 0.0001 between indicated groups. ##*p* < 0.01 between WT-CTRL and APP^NL-G-F^-CTRL or main-effect of PLX5622.

To directly determine if MGnD release more EVs compared to non-MGnD *in vivo*, we developed a novel microglia-specific lentivirus expressing mEmerald-CD9 (mE-CD9) fusion protein (Fig 4A) [36]. We replaced the promoter of the lentiviral vector with the EF1α promoter, which is highly active in microglia. Additionally, tandem miR-9 target sequence (miR9T) was incorporated into the 3’UTR to ensure silencing of gene expression in any non-microglial cells [37]. *In vitro* transduction into HEK293T cells demonstrates transgene expression in 48 hours (Fig S4A). mE-CD9 was enriched in the EV fraction of the conditioned media compared to the cell lysate (Fig. S4B). The high-titer (~ 1E+9 TU/ml) mE-CD9 lentivirus was bilaterally injected into the MEC of both WT and APP^NL-G-F^ mice at 6 months of age. Following 10-day incubation, microglia-specific expression of mE-CD9+ was detected by GFP antibody staining (Fig 4B) and enhanced in the mE-CD9^+^ microglia proximal to 82E1^+^ plaques. Double immunofluorescence of mE-CD9 and IBA1 shows that close to 100% of mE-CD9+ cells are IBA1+ microglia (Fig. S4C). MGnD were detected by Mac2 staining [38], which co-labeled MGnD with Clec7a^+^ nearly 100% of the time (Fig S4D) with a very small amount of MGnD not exhibiting Mac2 positivity. Z-stack images of Mac2^−^ microglia in WT mice, and Mac2^−/+^ microglia in APP^NL-G-F^ mice were captured via Leica SP8 with Lightning super-resolution confocal microscope and processed in IMARIS rendering software to swiftly and automatically quantify the number of mE-CD9^+^ particles surrounding individual microglia (Fig 4C). Quantification of mE-CD9^+^ particles localized around individual microglia revealed that Mac2^+^ MGnD released over 3 times as many EVs as Mac2^−^ microglia (Fig 4D). The intensity of Mac2 staining in microglia showed a strong positive correlation with the particle number of EVs, suggesting that there is a heightened release of EVs caused by the MGnD phenotype (Fig 4E). We collected the EV fraction of the left hemisphere of bilaterally-injected brains with discontinuous sucrose gradient ultracentrifugation as previously described [39, 40]. We performed mE-CD9 ELISA of whole brain homogenate and extracellular vesicles isolated from mE-CD9 lentivirus-injected brains, which showed a trend towards increased mE-CD9 concentration in APP^NL-G-F^ brains compared to WT brains (Fig 4F). We suspect this is due to the heterogeneous population of both Mac2+ and Mac2− microglia as a source of EVs in both groups. Taken together, these data demonstrate that Aβ plaque-associated MGnD secrete significantly more EVs than non-MGnD microglia in APP^NL-G-F^ mice in vivo.

**Figure 4.**
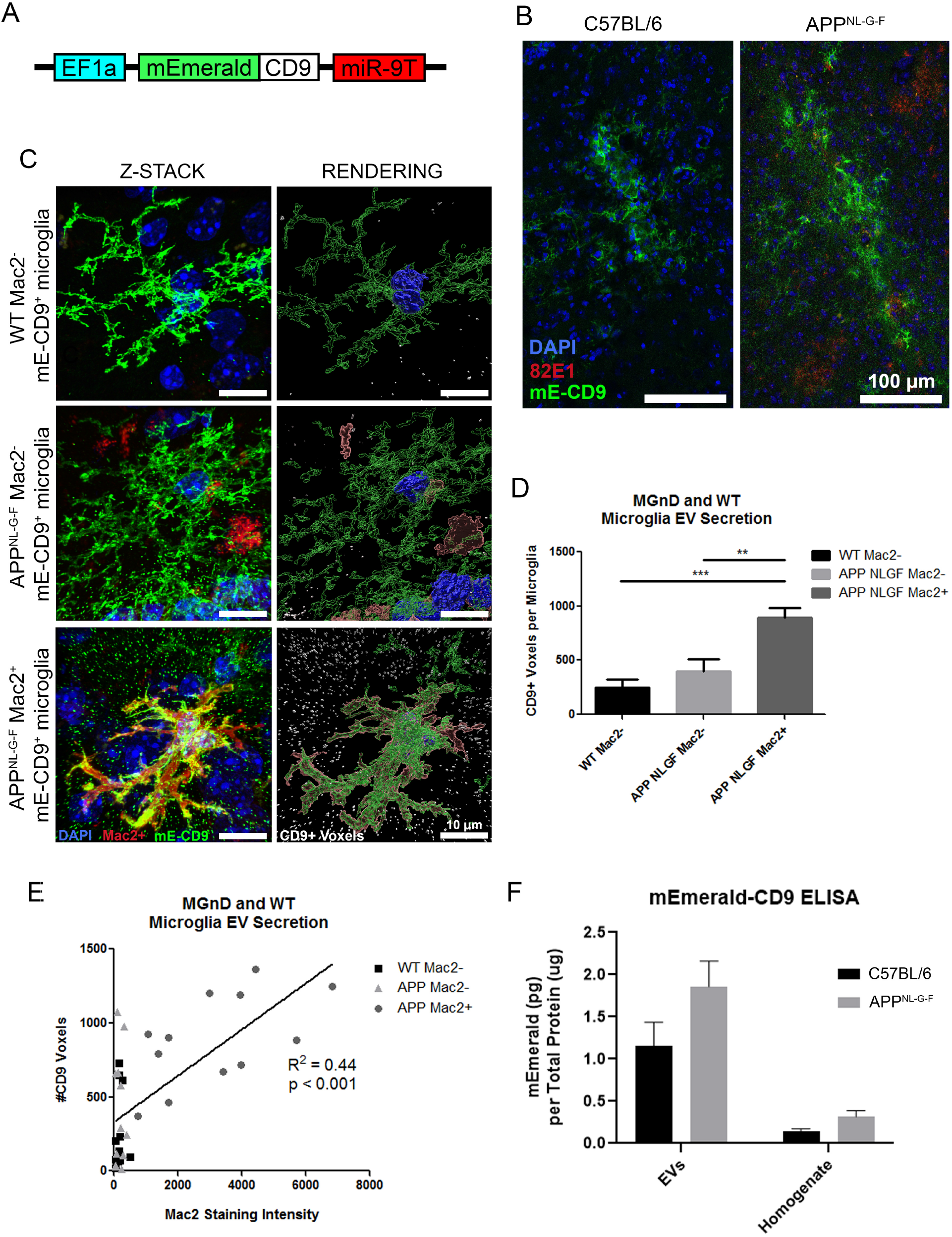
*In vivo* imaging of EV secretion from microglia after lentiviral microglia-specific expression of mE-CD9. **A.** Schematic of mE-CD9 lentiviral construct. **B.** Representative images of mE-CD9 lentivirus transduction in WT and APP^NL-G-F^ microglia following injection and 10 day incubation. **C.** Representative images of individual microglia and mE-CD9+ particles released. **D.** Unbiased quantification of mE-CD9+ voxels surrounding individual microglia (n = 12 microglia per group from 3 mice per group). **E.** Regression plot of Mac2 staining intensity versus the number of mE-CD9 particles released by individual microglia. **F.** Results of mE-CD9 ELISA from purified EVs and whole brain homogenate following 10-day incubation of mE-CD9 lentivirus. All values displayed in **A-F** represent the mean ± SEM for a minimum of 3 animals per group. Graphs comparing values across all 4 groups were analyzed via 2-way ANOVA with Tukey post-hoc analysis for individual comparisons. Graphs comparing two groups were analyzed via Unpaired t-test. ** *p* < 0.01, \*\*\**p* < 0.001, between indicated groups.

**Figure 5.**
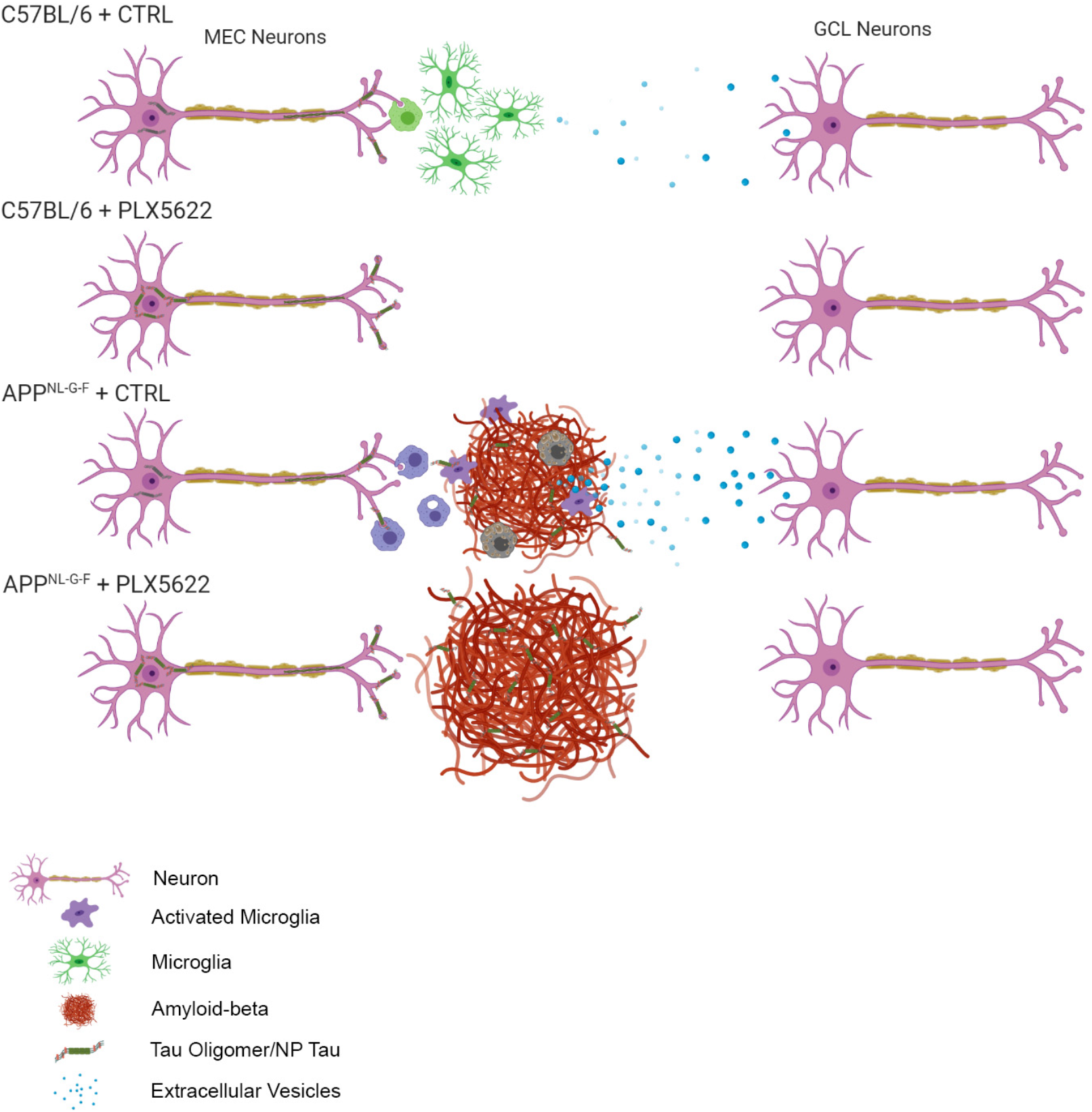
Schematic diagram of Aβ plaque deposition and microglia-mediated tau propagation. In WT mouse brain, tau propagation from MEC to GCL neurons is sensitive to microglia. In in APP^NL-G-F^ mouse brain, Aβ plaque associated microglia are more phagocytic and hyper-secrete tau-seeding EVs compared to homeostatic microglia, resulting in enhanced tau propagation from MEC to GCL regions. Both Aβ plaque and NP Tau pathology are increased following microglial depletion, suggesting their active roles on compaction and clearance of Aβ plaques and NP tau.

## Discussion

A number of publication including ours demonstrate the beneficial effect of CSF1R inhibitor-mediated microglial depletion on halting tauopathy in different tau mouse models [22, 41, 42]. One CSF1R inhibitor, JNJ-40346527, is being tested on the Phase 1 study of AD cases (NCT04121208). We previously established the role of microglia-derived extracellular vesicles in the propagation of pathologic tau from the MEC to the GCL of the DG [22]. In this study, we utilize the APP^NL-G-F^ mouse model to demonstrate how microglia react to amyloid pathology to become MGnD, which exacerbate propagation of tau through enhanced EV release. Treatment of mice with 1200 mg/kg PLX5622 resulted in ~92% reduction in the amount of microglia and ~80% of activated and plaque-associated microglia. It was noted previously that at lower doses of PLX5622 treatment, plaque-associated microglia are specifically depleted whereas homeostatic microglia are extant [19]. Another paper reported much less efficiency of microglial depletion by PLX3397 when administered to aged tau mice [23]. The general consensus of previous research regarding the effect of microglia depletion on amyloid deposition is that it is largely ineffectual aside from the 5xFAD mouse model, while we have found it causes a significant increase on amyloid deposition. There are several possibilities which may explain the discrepancy between ours and the previous findings. Firstly, all of the previous studies assessing the effect of microglia depletion on amyloid deposition used transgenic models that overexpress the amyloid precursor protein *APP* (Table S1). Deposition as a result of overexpression in combination with mutation may likely be more intense than the APP^NL-G-F^ model, which replaces the intrinsic m*APP* gene with the human version expressing three mutations to increase amyloid deposition [24]. Therefore, depletion of microglia in the former case may not have as large an effect on amyloid deposition as in the latter; overexpression of *APP* may saturate the ability of microglia to clear Aβ, resulting in no effect of depletion. Furthermore, the heterogeneity of mouse models, onset and duration of microglia depletion, and techniques for quantifying deposition all make comparisons difficult to interpret. Most studies in which amyloid burden is assessed utilize 6E10 for staining, which recognizes residues 1-16 of Aβ. In this study, we find no effect of microglia depletion on amyloid deposition using the 82E1 antibody, which recognizes the same residues. However, staining with 4G8, which recognizes residues 17-24 of Aβ, revealed a dramatic increase in diffuse plaque deposition that recapitulated the increases found on compact plaque deposition determined by Thioflavin S staining. These data suggest microglia are indeed involved in plaque compaction and clearance.

The role of microglia in mediating the interplay between amyloid and tau pathologies remains elusive. We previously reported the beneficial effects of microglia depletion as well as inhibition of EV synthesis in preventing tau spread in WT and PS19 mice [22, 43] We witnessed the same, but more enhanced effect in a mouse model exhibiting the other pathological hallmark of AD, amyloid plaques. In this study, overexpression of mutant P301L tau in the MEC of APP^NL-G-F^ mice produced two different kinds of tau pathology: increased AT8^+^ neurons at the GCL and NP tau on amyloid plaques. Aggregated and phosphorylated tau in dystrophic neurites, recently described as “NP Tau”, is a known pathology of AD brains [10, 13]. We observed an increase in NP tau accumulation by over-expressing P301L tau mutation in the MEC of APP^NL-G-F^ mice. This is supported by the recent study showing that the tau fibril injection in APP^NL-G-F^ mice induced NP tau accumulation [10]. We found that microglia depletion increased NP tau accumulation suggesting that plaque associated microglia may actively phagocytose NP tau. This is consistent with the recent report demonstrating the enhanced NP tau formation in APPPS1-12 mice in TREM2 knockout or R47H mutant background, which show diminished number of plaque-associated microglia [13] Authors speculated that microglial dysfunction increases the susceptibility of dystrophic neurites to develop this pathology, possibly through increased vulnerability to the toxic effects of Aβ42 in plaques. Our study provides an additional interpretation: NP tau may be actively phagocytosed by plaque-associated MGnD and their depletion enhances NP tau pathology. The phagocytosed NP tau may also be an important source of microglia to secrete tau seed-containing EVs.

Axons projected from the MEC through perforant path form synapses with distal dendrites from GCL neurons in the OML, allowing for pathological tau seeds to spread from neurons to neurons [31, 44, 45]. It is well known that microglia actively phagocytose synapses by synaptic tagging with complements (C1q and C3) [46, 47], which also play an important role for Aβ-induced synaptic loss [48]. As shown in the current and previous studies [34, 49], C1q signal is highly intense in the OML, and this study shows that it is microglia-dependent. C1q is likely to play a role in engulfing damaged synapses containing pathological tau seeds by microglia in the OML. Given that microglia are known to be activated and recruited to the plaque region where they surround, phagocytose, and also secrete toxic aggregates through their EVs, we hypothesize that the activity of plaque-associated microglia in this region may inadvertently bolster tau propagation from the MEC to the DG in APP^NL-G-F^ mice. Indeed, microglia-derived EVs are endocytosed by proximal neurons and influence their activity [50–52]. Another recent report shows that microglia isolated from AD brain or tau mice can seed tau *in vitro* [53]. Interestingly, we found that microglia depletion dramatically reduced the amount of EVs in the plaque region, suggesting that activated, plaque-associated microglia hypersecrete EVs and may increase the rate of tau propagation through EVs. Indeed, many of the EV markers, such as CD63, CD9, CD81, and Tsg101 are upregulated in DAM/MGnD microglia isolated from APP or APP/PS1 mice [8, 54], a part of which was recapitulated using isolated Clec7A^+^ MGnD microglia and microglia depletion-sensitive co-localization of Tsg101 and Clec7A around Aβ plaques in this study.

Given that immunofluorescence staining of tetraspanins and other EV markers is cumbersome and not guaranteed to be reflective of EV release from microglia, we sought to develop a way to measure EV release specifically from microglia *in vivo*. We accomplished this by constructing the mE-CD9 lentivirus which robustly transduces and expresses fluorescent mEmerald conjugated to CD9 in microglia. This allowed us to easily quantify the number of mE-CD9+ particles localized around infected microglia. We found that EV release appeared to be over 3 times higher from MGnD compared to homeostatic microglia and that the more microglia exhibit the MGnD phenotype, the more robust the EV release. Coupled with the qPCR and Tsg101 staining results, there is strong evidence to suggest the MGnD phenotype enhances EV release *in vivo*.

In summary, we show that amyloid plaques enhance the propagation of pathologic tau by induction of plaque-associated MGnD, which are hyper-phagocytic and hyper-secreting EVs. We also show that in APP^NL-G-F^ mice, compaction, size and the number of Aβ plaques are sensitive to the microglial depletion, and that AAV-P301Ltau injection enhances NP tau, which became more prominent by microglial depletion. We also shows that the OML region is particularly tagged by microglia-derived C1q, indicative of active niche for microglial synaptic pruning. Therefore, microglia are likely exacerbating the spread of pathologic tau through EV secretion, suggesting a possible new therapeutic target for preventing the progression of AD pathology.

## Methods

### Animals and Genotyping

All mouse care and experimental procedures were approved by Institutional Animal Care and Use Committee of the Boston University School of Medicine. APP^NL-G-F^ mice were bred and genotyped in-house. C57BL/6 were purchased from the NIA. Mice were caged in accordance with their own sex and housed in a barrier facility with 12 hours light and 12 hours dark cycles. Roughly half of the animals used were male and half female between WT and APP^NL-G-F^. Food and water was provided *ad libitum*. Throughout the life of all mice, veterinary staff closely monitored animals for complications. Genotyping for animals was conducted in house via PCR using the following primers [24]: E16WT: 5’ – ATCTCGGAAGTGAAGATG – 3’ E16MT: ATCTCGGAAGTGAATCTA WT: 5’ – TGTAGATGAGAACTTAAC – 3’ loxP: 5’ – CGTATAATGTATGCTATACGAAG – 3’

### Viral vector production

The AAV-SYN1-P301L tau (AAV-P301Ltau) was generated as previously described [22]. This virus is recombinant AAV pseudoserotype 2/6 (AAV2/6) and was created via transient transfection in human embryonic kidney (HEK293) cells. The vector consists of AAV-2 inverted terminal repeats, the human synapsin-1 gene promoter driving expression of human P301L tau 1-441, the woodchuck hepatitis virus post-transcriptional control element (WPRE) and a bovine growth hormone polyadenylation site. Iodixanol step gradient ultracentrifugation followed by heparin FPLC affinity chromatography and dialysis in PBS overnight was used to purify and prepare AAV particles. Viral titers were calculated from Q-PCR. Purity of AAV was determined by SDS-PAGE and coomassie brilliant blue staining.

### CSF1R inhibitor treatment and Intracranial Injection

PLX5622 or control (Plexxikon, Inc., San Francisco, CA), was impregnated into rodent chow at 1200 mg/kg (AIN-76A, Research Diet, Inc., Brunswick, NJ) and provided the animals in equal quantity starting at 4 months of age. After one month since the beginning of PLX5622 treatment, intracranial injections of AAV-P301Ltau were administered to the drug-treated mice. This AAV expresses the mutant version of human tau P301L under the syn-1 promoter, which is neuron-specific [55]. Nine tenths of 1 μl were injected into each mouse with coordinates (AP: 4.75, ML: 2.90, DV: 4.64) at a viral titer of 1.2 × 10^11^ using a Hamilton model 701 LT 10 μl syringe (#80301) and Neurostar glass capillary held together with microelectrode holder (World Precision Instruments #MPH6S10). Mice were anesthetized during the procedure with 3% isoflurane and received 15 mg/kg Meloxicam for pain relief. Experimenters were blinded to which group received PLX5622 and which received placebo throughout the experiment. Animals experiencing undue trauma or distress following injections were sacrificed and excluded from the study. Following one-month incubation of virus with continuous PLX5622 treatment, animals were sacrificed with transcardial perfusion with PBS and followed with fixation with 4% paraformaldehyde (PFA) solution. Brains were immediately harvested.

### Histological Processing and Immunofluorescence staining

Brains were immersed in 4% PFA at 4 degrees Celsius for overnight following harvest. The following day, brains were placed in 30% sucrose solution in PBS in 4 degrees Celsius for overnight again in preparation for cryosectioning. Sagittal brain slices were then gathered using a Thermo Fisher Scientific Cryostar NX50 (# 957250K) at 30 μm each. Following sectioning, brains were mounted on superfrost plus microscope slides (#22-037-246, Thermo Fisher Scientific) and stored at −80 degrees Celsius. The following antibodies and reagents were used for immunofluorescence staining: 4G8 1:100 (BioLegend #800704), AT8 1:300 (Thermo #MN1000), GFAP 1:300 (Cell Signaling Bio #36705), C1Q 1:300 (Abcam #ab182451), 82E1 1:100 (IBL #10323), Thioflavine S 1:50 (Sigma #T1892-25G), HT7 1:300 (Thermo #MN1000), Clec7A 1:100 (InvivoGen #mabg-mdect), Tsg101 1:300 (Santa Cruz #sc-7964), DAPI 1:2500 (Thermo #62248). Rabbit polyclonal anti-mouse P2RY12 antibody 1:600 was generously provided by Oleg Butovsky [56, 57]. Sections were washed with PBS prior to antigen retrieval with 88% Formic Acid. Blocking solution consisted of 5% normal goat or donkey serum, 5% bovine serum albumin (BSA), and 1% Triton X-100. All following primary and secondary staining buffer consisted of 5% BSA, 1% Triton X-100 in PBS. Following staining, sections were allowed to air-dry before coverslips were mounted using Fluoromount G (Invitrogen #00-4958-02). Tiled images were taken at 20X with a Nikon Eclipse Ti microscope. Confocal images were taken using Leica SP8 Confocal Microscope with Lightning. Images were observed and analyzed using open source image processing package FIJI. Processing of z-stack images and quantification of mE-CD9^+^ particles was accomplished using IMARIS rendering software.

### Lentiviral vector production

The pLV-EF1α-mEmerald-CD9-miR9T lentivirus was generated by modifying commercially available pLV.PGK.GFP.miR9T lentivirus [37]. The vector backbone was modified to contain the murine EF1α promoter and express mEmerald conjugated to CD9 followed by miR9T in the 3’UTR. Viral particle production and packaging was done by a commercial source (SignaGen Laboratories, Rockville MD, USA). The viral titer is > 1E + 9 TU/ml. Lentivirus (1 μl) was injected to the MEC (AP: 4.75, ML: 2.90, DV: 4.64) of 6 month-old APP^NL-G-F^ and C57BL/6 mice, and euthanized at 10-day post-injection for immunohistochemical and biochemical analyses.

### Statistical Analysis

All statistical analyses were performed in GraphPad Prism v8 (Graph-Pad Software, Inc). Twoway ANOVA was used to assess comparisons between all four experimental groups when applicable. Normal distributions were assumed when making post-hoc analyses and correcting for multiple comparisons (Tukey). In instances of two group comparisons, unpaired t-tests assuming equal variances were used.

## Abbreviations

AAV: adeno-associated virus
Aβ: amyloid-β peptide
AD: Alzheimer’s disease
APOE: Apolipoprotein E
CA1: Cornus Ammonis 1
Clec7A: C-type lectin domain family 7 member A
*CSF1R*: Colony stimulating factor 1 receptor
DG: Dentate Gyrus
EVs: Extracellular vesicles
GCL: granule cell layer
GFAP: Glial fibrillary acidic protein
HEK293: Human embryonic kidney cells
mE-CD9: mEmerald-CD9 conjugate
MGnD: Disease-associated neurodegenerative microglia
MEC: Medial Entorhinal Cortex
mi9RT: tandem miR-9 target sequence
NFTs: neurofibrillary tangles
OML: outer molecular layer
PFA: paraformaldehyde
pTau: phosphorylated-Tau
TA: Temporoammonic
TREM2: Triggering Receptor Expressed on Myeloid Cells 2
Tsg101: Tumor susceptibility gene 101
WT: wild type

## Acknowledgments

The authors thank Jeremy Kwon, Abigail Tesfaye and Kimberly Soto for their technical assistance in image analyses and Dr. Zhi Ruan for suggestion for NP tau characterization. The PLX5622 compound was obtained from the company PLX. The Neurostar technical support team was also very helpful in establishing the brain injection system.

## Availability of data and materials

Data produced and analyzed in this study are all available herein. Datasets used for analysis are available from the corresponding author upon reasonable request.

## Authors’ contributions

KC performed stereotaxic injections, immunohistochemistry, and image analyses. JD trained KC on how to perform surgeries and immunofluorescence, aided in statistical analyses, and contributed to manuscript preparation. SH and NI designed and constructed lentiviral vectors and performed in vitro experiments. SI bred and provided all mice required for the study and contributed to manuscript preparation. TS and TCS provided APPNL-G-F mice, technical suggestions for immunohistochemistry and edited manuscript. TI designed the study and contributed to supervision, manuscript preparation and editing. All authors have read and approved the final version of the manuscript.

## Consent for publication

All involved parties consented to publication of this work.

## Ethical approval and consent to participate

Animals and experimental procedures were approved by the IACUC of Boston University.

---

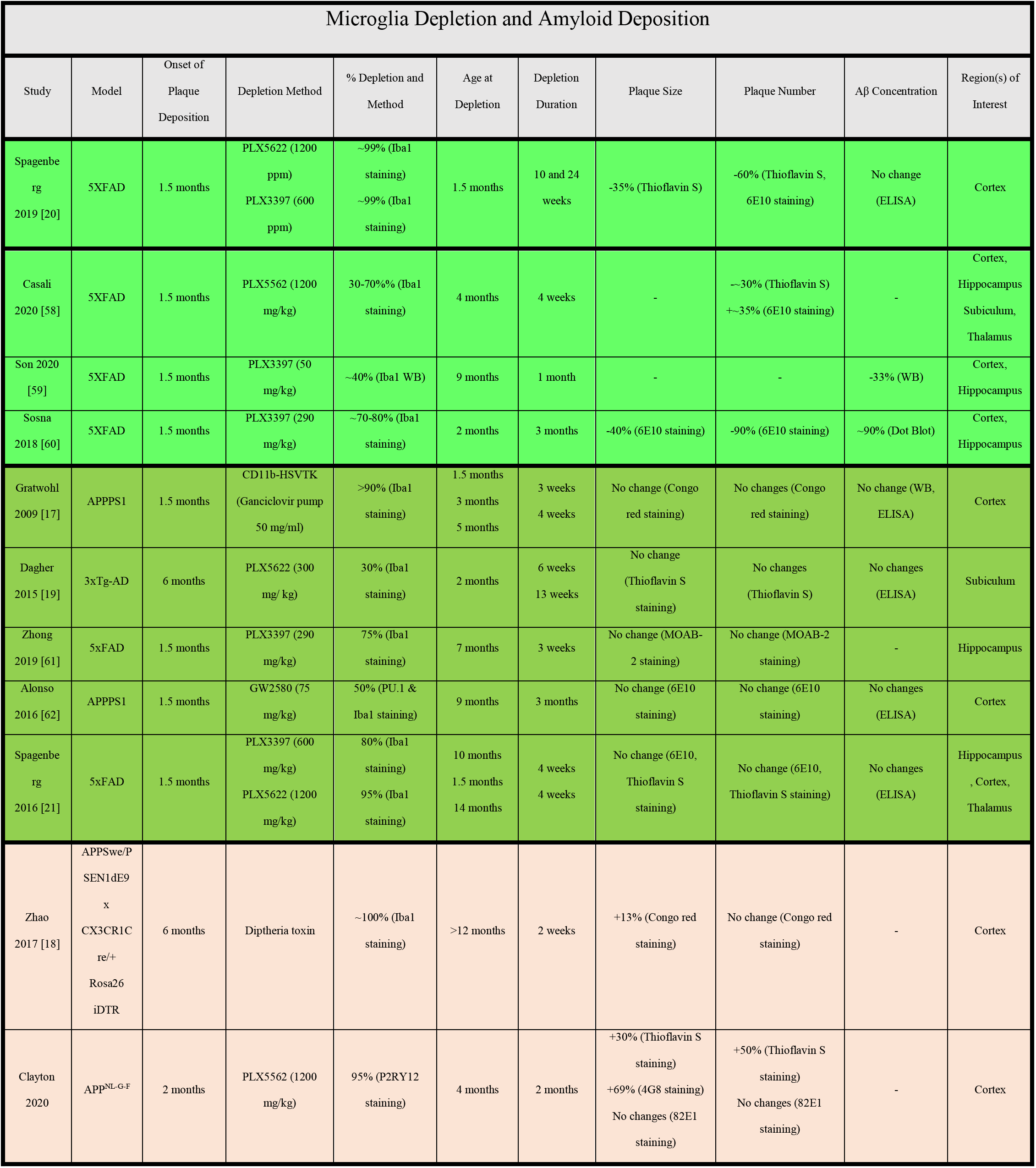

**Fig. S1.**
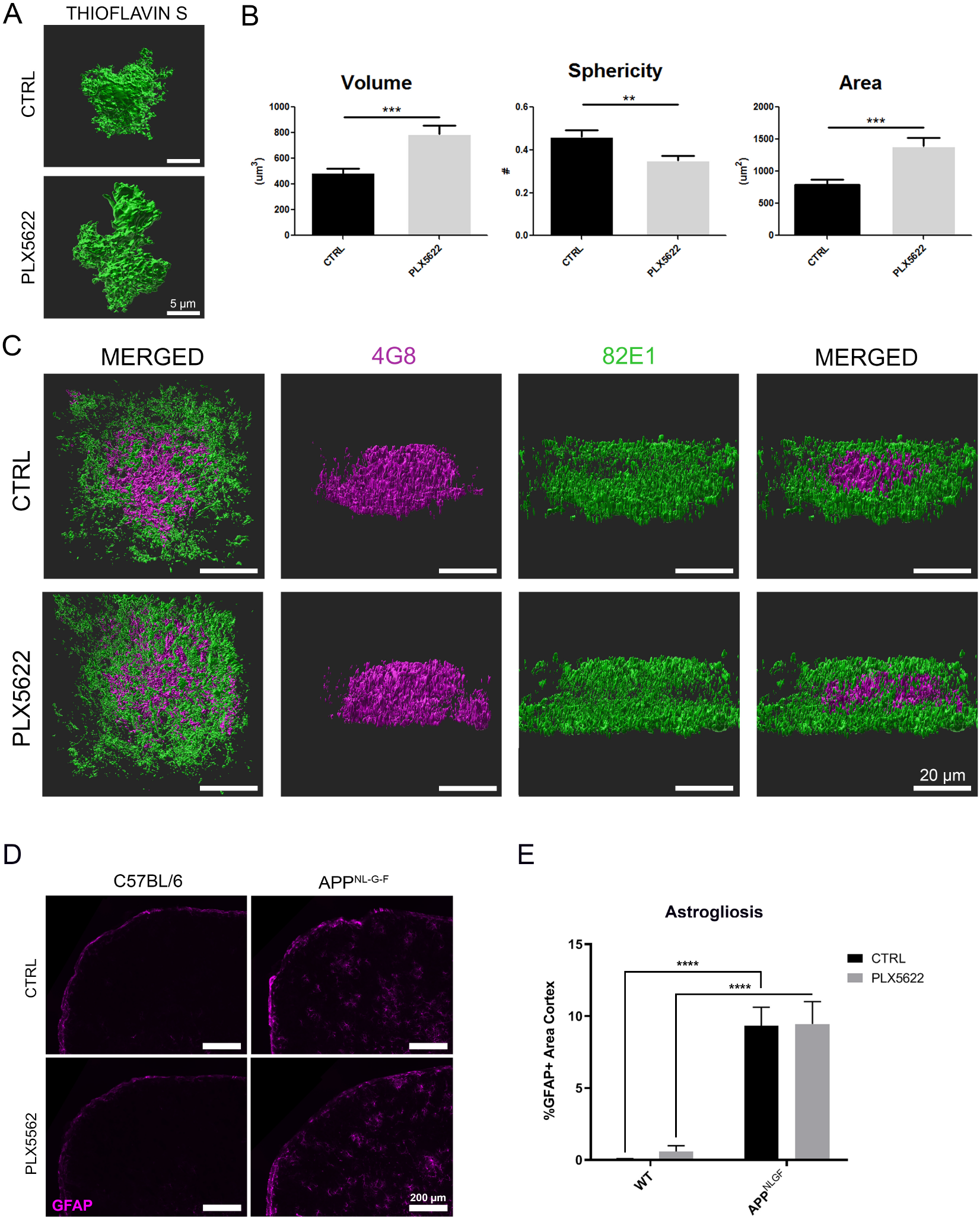
Effects of microglia depletion on plaque deposition and astrogliosis. **A** Representative 3D rendering of Thioflavin S^+^ plaques with and without microglia depletion. **B** Unbiased quantification of plaque volume, sphericity, and overall area (n = 20 plaques per group). **C** Representative 3D rendering of 4G8^+^ and 82E1^+^ plaques showing distinct staining patterns. **D** Representative images of GFAP staining in the cortex. **E** Unbiased quantification of GFAP^+^ area in the cortex. All values displayed in **B** represent the mean ± SEM for a minimum of 4 animals per group. Graphs comparing values across all 4 groups were analyzed via 2-way analysis of variance (ANOVA) with Tukey post-hoc analysis for individual comparisons. ** *p* < 0.01, *** *p* < 0.001, **** *p* < 0.0001.

**Fig. S2.**
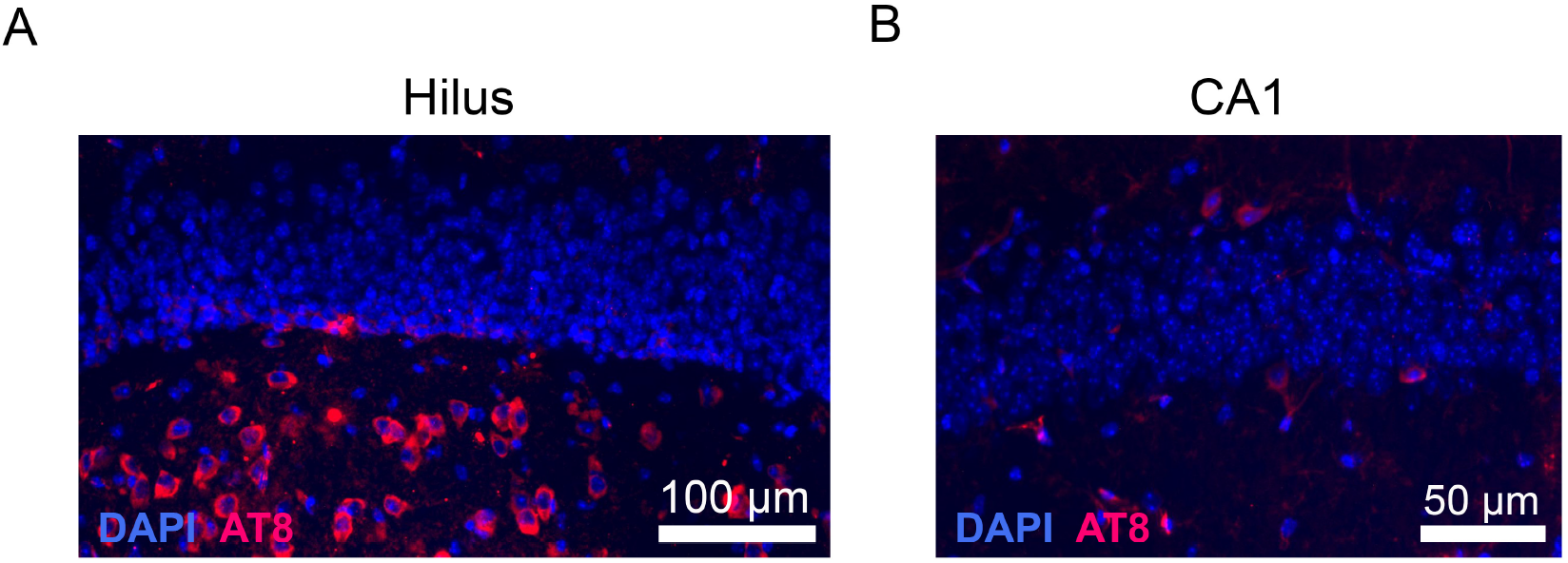
**A** Representative picture of strong tau propagation to the Hilus region of the hippocampus. **B** Representative image of tau propagation to the CA1 region of the hippocampus.

**Fig. S3:**
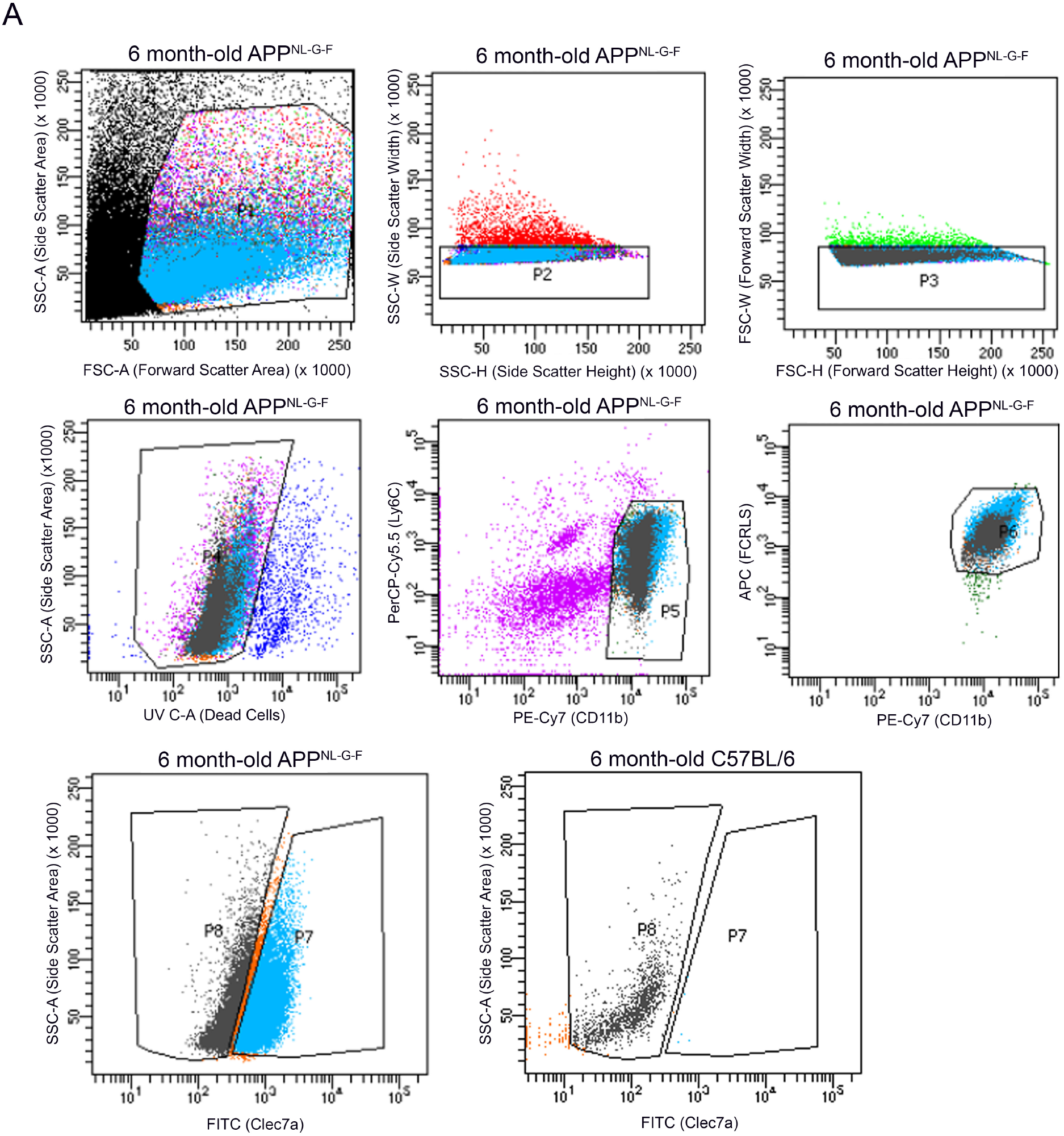
**A** Gating settings for fluorescence activated cell sorting of Cd11b^+^ / Ly6c^−^/ Fcrls^+^ / Clec7a± microglia from the brains of 6 months-old APP^NL-G-F^ mice. Following live/dead cell and singlet selection, microglia were selected via a combination of Ly6C, CD11b, and FCLRS prior to being sorted as Clec7a^+^ or Clec7a^−^.

**Fig. S4:**
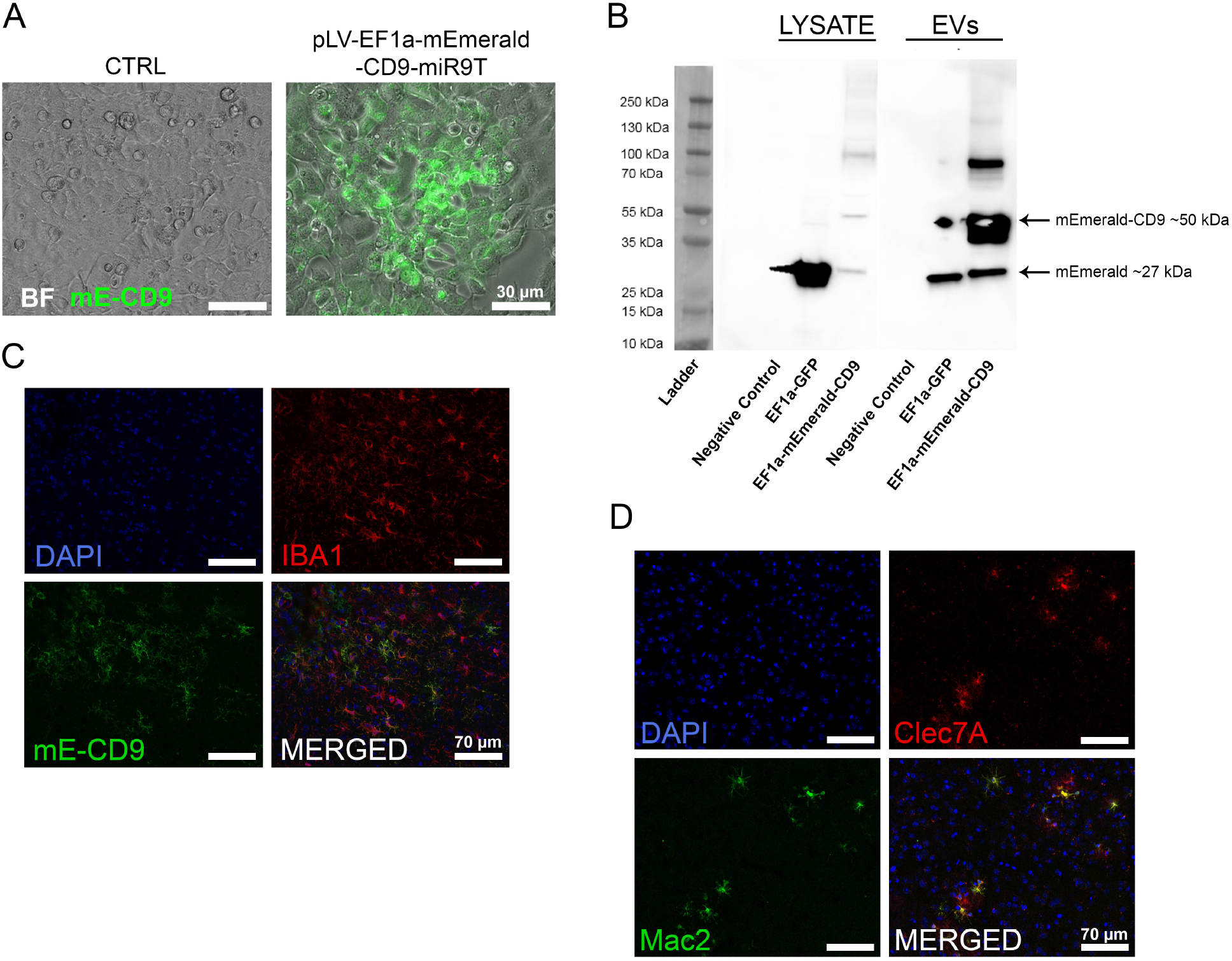
**A** Representative images of mEmerald-CD9 (mE-CD9) expression following control lentivirus (left) and mE-CD9 lentivirus (right) transduction in HEK293 cells following 48 h incubation. **B** Western blot of cell lysates and isolated EVs from transiently-transfected HEK293 cells demonstrating expression of mE-CD9. **C** Representative image of 10-day lentivirus transduction into the MEC, demonstrating ~100% colocalization of mE-CD9 with IBA1. **D** Representative images of overlap between Clec7A and Mac2 staining, demonstrating that of ~100% Mac2^+^ cells are Clec7A^+^ MGnD microglia.

